# The Utility of Fibroblast Co-culture for the Maintenance of Episomes in Human Papillomavirus-Associated Cancer Models

**DOI:** 10.1101/2025.10.06.680765

**Authors:** Austin J. Witt, Rachel L. Lewis, Xu Wang, Phoebe Bridy, Jenny Roe, Iain M. Morgan, Claire D. James, Molly L. Bristol

## Abstract

Human papillomavirus-associated head and neck squamous cell carcinomas (HPV+ HNSCCs) lack early diagnostics and continue to rise in incidence. HPV16 has been detected in ∼90% of HPV+ oropharyngeal cancers (HPV+ OPCs), an anatomical subset of HNSCC, with the majority retaining episomal viral genomes. Despite this, existing episomal HPV16+ OPC cell lines are especially scarce. UMSCC104s were initially reported as episomal; however, the literature contains conflicting reports regarding the genome status of these cells. We now show that UMSCC104s rapidly undergo integration and E2 loss under standard monoculture, and later lots are fully integrated. These findings resolve prior discrepancies and underscore the instability of episomes in monoculture. Accurate models of HPV-driven cancers are critically needed. We propose fibroblast co-culture methods, traditionally utilized for HPV+ keratinocyte models, as a strategy to preserve episomal status in cancer models by supporting viral and host genome stability.

## Introduction

Human papillomaviruses (HPVs) are oncogenic viruses responsible for approximately 5% of human cancers, including a growing subset of head and neck squamous cell carcinomas (HNSCCs)^1–3^. Among these, HPV+ oropharyngeal cancers (HPV+ OPCs) have risen sharply in incidence, with HPV16 detected in as high as 90% of cases^3,4^. While HPV+ OPC patients generally have favorable prognoses, the substantial long-term morbidities among survivors highlight the urgent need for improved therapeutic approaches^4–6^. To this point, there is a significant need for appropriate preclinical models to study this disease.

HPV episomal genome status is prognostically significant in HNSCC, and more than 70% of HPV+ OPC patients retain extrachromosomal viral episomes^6–9^. Episomal status is associated with significantly improved overall survival, when compared to tumors with integrated genomes or HPV negative disease^6^. Conversely, most existing HPV16+ HNSCC cell lines are derived from non-oropharyngeal sites and contain integrated viral genomes, thus limiting their clinical relevance^3,6–11^. The successful establishment of HNSCC cell lines has been shown to correlate with poor prognosis for the associated patients^9,12^. This may partially account for the selective availability of cell lines harboring integrated viral genomes, as they have poor prognostics and in turn may be more likely to yield stable cell lines than those retaining episomal HPV. We also suggest that the traditional monoculture approach commonly utilized for cancer cells may likewise hinder the establishment of episomal lines, consistent with knowledge that fibroblast co-culture is essential for episomal maintenance in other epithelial models^13^. This idea is further supported by our recent observation that 50% of a set of HPV16+OPC patient derived xenografts (PDXs) retain episomal genomes, potentially due to their consistent maintenance adjacent to mouse stroma^3^.

Upon initial HPV infection, viral genomes are maintained as low-copy extrachromosomal episomes in basal keratinocytes^13^. *In vivo*, infected basal keratinocytes are adjacent to the stroma, and fibroblasts are a major component in this microenvirnoment^14^. Fibroblasts are noted to secrete essential growth factors that regulate both keratinocyte proliferation and differentiation^10,13^. *In vitro*, episomes are generally unstable unless HPV+ keratinocyte cell lines are established and maintained in a co-culture system utilizing inactivated 3T3-J2 murine fibroblasts (J2s) ^10,13,14^. It is well established that without J2s, HPV+ keratinocyte cultures rapidly integrate the viral genome^10,13^. Of note, cervical intraepithelial neoplasia (CIN) episomal cell lines, such as W12 (HPV16) and CIN612-9E (HPV31), were also established using fibroblast co-culture conditions^15^. As feeder-layer fibroblasts are vital for establishing and maintaining episomal keratinocyte and CIN models, we further suggest that J2 co-culture methods should be adapted and utilized for the establishment and maintenance of episomes in clinically relevant HPV-associated cancer models^9,10^.

## Results

### Rapid Episome Loss in UMSCC104s When Grown in Traditional Monoculture

Only a limited number of HPV+ HNSCC cell lines are commercially available, with most harboring integrated viral genomes; this does not reflect the clinical abundance of episomal HPV+ OPC^6,7,9^. UMSCC104s (HPV16+ HNSCC cell line) were originally reported to contain episomal HPV16 genomes^16^. Of note, there is significant disagreement in the literature as to the episomal status of these cells^11,16,17^. We investigated whether this may be due to integration of the viral genome during passaging. Passage 3 UMSCC104 cells, from a lot originally obtained in 2015, demonstrate the presence of episomal DNA via Southern blotting (Figure 1A – for methodology see^18^). These were compared against UMSCC47 and UMSCC152 cells, containing known integration sites, as well as HPV-negative HN4 cells^11,17,19^. The ExoV exonuclease assay provides a more sensitive and quantitative method for detecting HPV16 episomes and genome integration events, offering distinct advantages over Southern blotting for tracking integration dynamics^3,19^. Using the ExoV exonuclease assay, we observed that viral genome integration occurs rapidly during passaging of UMSCC104 (Figure 1B, for methodology see references)^3,19^. Within 25 passages, UMSCC104 viral genome status transitioned from primarily episomal to fully integrated, and both E2 and E6 viral DNA presence was significantly reduced (Figure 1B); of note E2 and E6 DNA levels remained detectible over background. Moreover, UMSCC104 lots obtained in 2024 were fully integrated upon receipt when measured by ExoV (Figure 1C), underscoring the inherent instability of episomes under basic vendor monoculture maintenance and expansion protocols prior to distribution.

**Figure 1.**
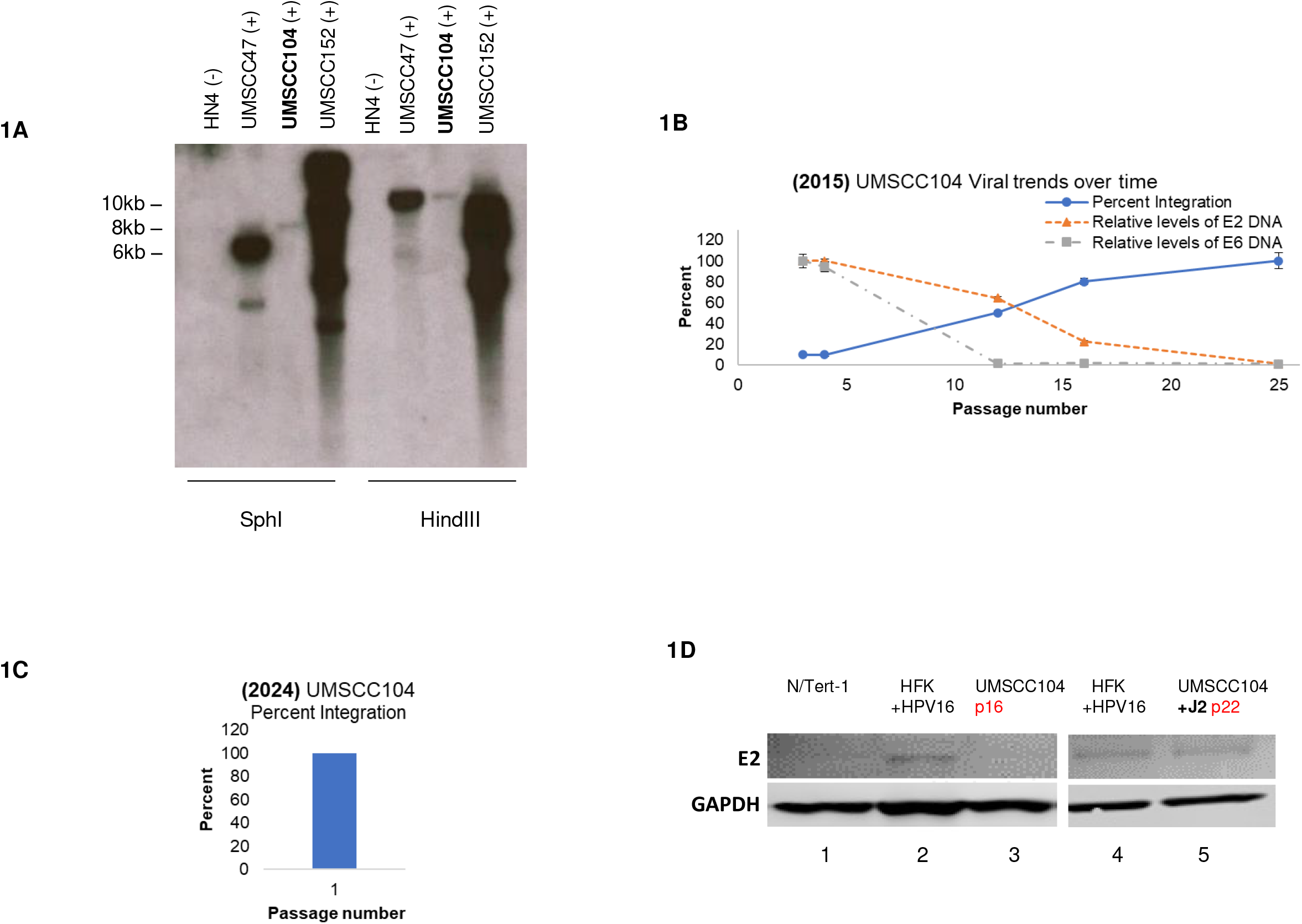
HPV episomal instability of UMSCC104 cells grown with traditional monoculture methods. **1A**. Total cellular DNA was extracted from HN4, UMSCC47, UMSCC104, and UMSCC152 cell lines. 5µg of DNA was digested with either SphI (to linearize the HPV16 genome) or HindIII (which fails to cut HPV16 genome). Membranes were probes with radiolabeled (32-P) HPV16 genome and then visualized by exposure to film. Image presented was after 1 hour exposure. **1B**. Early passage UMSCC104s, obtained from Millipore in 2015, were serially passaged and analyzed via the ExoV exonuclease assay at noted passage numbers (x-axis). Percent integration was calculated from the ratio of circular and linear E2 viral DNA in comparison to host mitochondrial (circular) and GAPDH (linear). Relative observed DNA levels of HPV16 E2 and E6 are also presented at these time points. **1C**. Passage 1 UMSCC104s, obtained in 2024 from Millipore, were analyzed via the ExoV exonuclease assay. Percent integration was calculated as noted in 1B. **1D**. Passage 12 UMSCC104 cells were sequentially passaged in the presence or absence of J2. Western blotting for E2 was compared to E2 ± keratinocyte cell lines at the noted passage numbers.

### Fibroblast Co-Culture Preserves Episomal Status of UMSCC104s

Fibroblast co-culture has been well established as a method to preserve episomal status of HPV+ epithelial cultures^10,13^. With that knowledge, we adapted this co-culture approach to determine if fibroblasts would also allow for the support and maintenance of the viral genome in UMSCC104s. Passage 12 UMSCC104 cells, have a proportionate ratio of integrated and episomal genomes, as demonstrated in Figure 1B. These cells were then grown in the presence or absence J2s. As E2 expression is known to correlate with episomal status, E2 protein expression was monitored in UMSCC104 and compared to cells with known E2± status (Figure 1D). UMSCC104s, grown in the traditional monoculture for four passages, had little-to-no detectible E2 protein expression (lane 3), when compared to HPV negative N/Tert-1 (lane 1) and episomal HPV16+ human foreskin keratinocytes (HFK+ HPV16, lane 2) consistently maintained on J2. Conversely, UMSCC104 cells maintained on J2 for eight passages had similar E2 protein expression levels as HFK+ HPV16 (lanes 5 and 6, respectively). At present, fibroblast coculture methods have demonstrated stable maintenance of episomal HPV16 genomes and E2 protein expression across >30 passages (not shown).

### Fibroblasts Support Viral and Host Genome Stability to Reduce Overall Integration Risks

We previously demonstrated that fibroblasts regulate the transcriptional signature of HPV+ keratinocytes^10^. In our RNA sequencing (RNAseq) analysis, we observed that the removal of fibroblasts significantly reduced the gene expression of numerous DNA repair pathways^10^. With that knowledge, we wanted to determine the effect of fibroblasts on DNA integrity in our various HPV+ models. N/Tert-1 cells (hTERT immortalized human foreskin keratinocytes), N/Tert-1 cells stably maintaining HPV16 episomal genomes (N/Tert-1+HPV16), HFK+HPV16 (foreskin keratinocytes immortalized by the entire HPV16 genome, replicating as an episome), and passage 14 UMSCC104s were grown in the presence or absence of J2s for one week^10,20^. The comet assay is a standard approach for measuring DNA strand breaks in single cells via electrophoresis^21^. In the comet assay, intact supercoiled DNA remains within the dense ‘head’ of the comet, whereas DNA fragments generated by strand breaks migrate and form a diffuse ‘tail’^21^. The extent of DNA damage is then quantified by the olive tail moment, which is a measure of the tail length and the fraction of total DNA present in the tail^21^. Fibroblast co-culture supported DNA integrity in all HPV+ cell lines, as observed by significantly smaller olive tail moments in Figure 2A (for methodology, see^22^).

**Figure 2.**
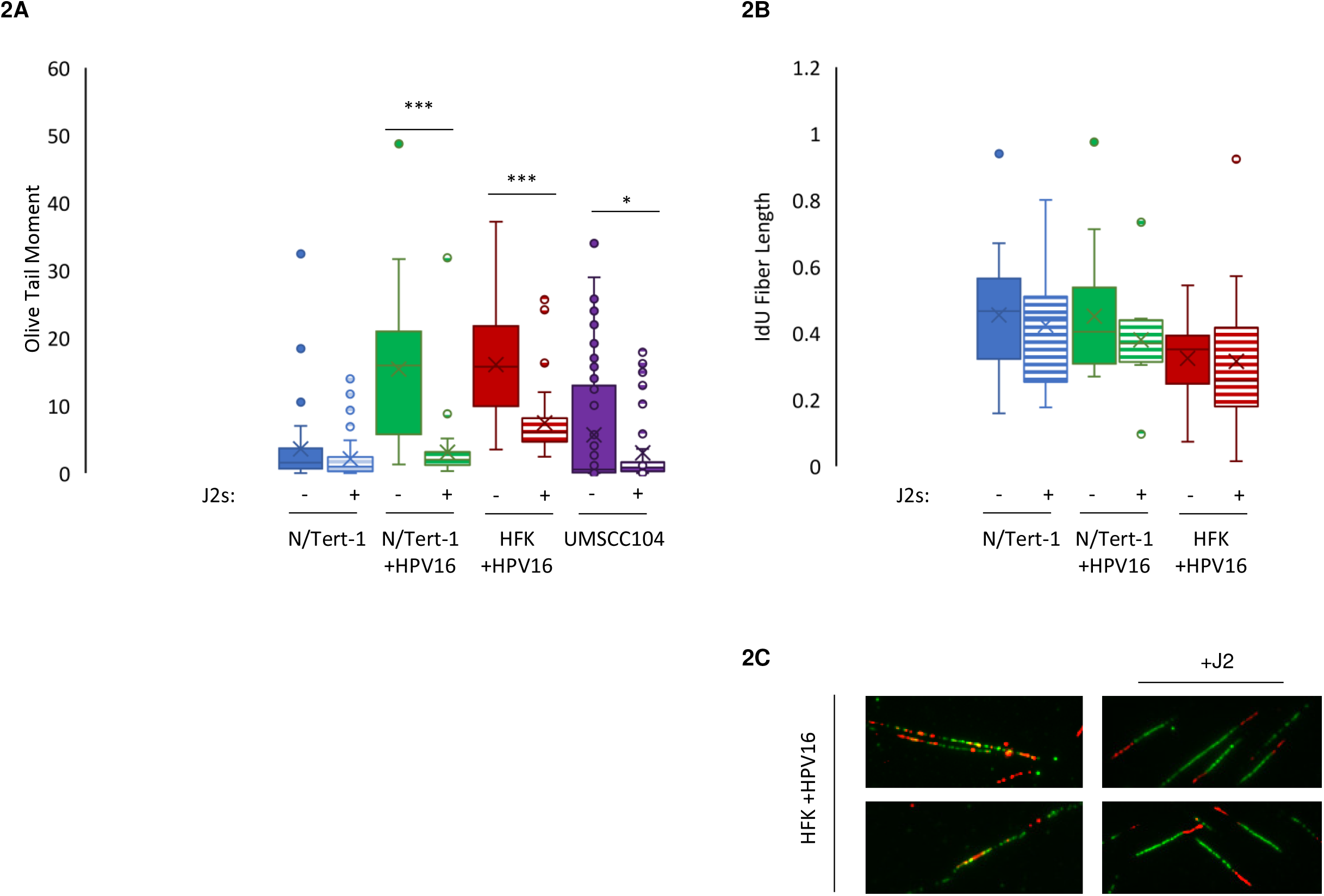
Fibroblasts enhance host genome stability, reducing the risk of viral integration. Noted cell lines were grown in the presence or absence of J2s for one week. UMSCC104 cells utilized were passage 14 from the Millipore lot obtained in 2015. **2A**. DNA strand breaks were assessed using the single-cell gel electrophoresis (COMET) assay. Cells were grown in 24 h then trypsinized, washed, resuspended in 0.5% low molecular weight agarose, and subjected to single cell gel electrophoresis. DNA was stained with SYBR Gold. Five randomly selected fields were imaged at 20× per replicate per cell line. The olive-tail moments (OTMs) of all nonoverlapping comets in each high-power field were quantified using CaspLab COMET assay software. Average OTM +/- SEM. **2B**. Cells were grown for 48 h and fiber assays were carried out in indicated cell lines. A summary of <green= IdU fiber length is shown. The length of labelled tracks was measured using ImageJ software. A minimum of 100 individual fibers were analyzed for each experiment and each experiment was repeated two times. **2C**. Representative <aberrant= fibers observed in HFK+HPV16 cells grown in the absence of J2 are shown in comparison the representative <normal= fibers observed in these cells maintained in fibroblast co-culture. Similar aberrant fibers were observed in N/Tert-1+HPV16 cells (not shown).

As the total level of observable DNA damage in a comet can be influenced by rates of repair, we also assessed replication fork dynamics utilizing the DNA fiber assay^23^. Removal of fibroblasts resulted in significantly longer IdU fiber portions in all of our HPV+ keratinocytes, indicative of faster replication speeds (Figure 2B, for methodology see^24^). HPV+ keratinocytes grown in the absence of fibroblasts also had numerous instances of multiple replication origins per fiber and fork termination events, suggesting aberrant repair dynamics (representative images in Figure 2C)^23^. Future analysis will utilize HU labeling to further investigate fork stability, fork initiation, fork stalling, and fork degradation. Of note, despite repeated attempts, we were unable to CldU label UMSCC104 cells. We therefore refrain from drawing definitive conclusions about fork dynamics of these cells at this time. Overall, the combined increase in DNA damage (Figure 2A) and altered replication dynamics (Figure 2B) in the absence of fibroblasts would enhance opportunities for viral integration events.

## Discussion

Our findings demonstrate that conventional cancer cell line monoculture approaches do not support episomal maintenance in HPV+ HNSCC lines; without stromal support, viral integration and altered viral genome regulation occurs in cell lines that were episomal when derived from patients. Episomal instability undermines the development of representative cell lines for studying HPV cancer biology and appropriate therapeutic responses. In contrast, fibroblast co-culture methods supported episomal stability in UMSCC104s, paralleling prior successes in HPV+ keratinocyte and CIN models. This strategy holds significant promise for generating more representative HPV+ OPC models. Episomal lines would better recapitulate the clinical presentation of HPV+ OPC. Of note, when UMSCC104s were retrieved from cryopreservation on fibroblasts, they have substantially improved recovery rates (not shown). We suggest that fibroblast co-culture may increase the overall success rate of establishing HPV+ lines from patient samples, a longstanding challenge in the field. While episomal cell lines are lacking, it is worth noting that episomal PDXs have been established^3^. While episomal HPV16+ OPC cell lines are lacking, 50% of HPV16+ OPC PDXs retain episomes^3^. The PDX episomal ratio much more closely reflects the ∼70% episomal prevalence in patients^6–8^. We further suggest that the more successful establishment and maintenance of episomal HPV16+ OPC PDXs may be supported by the adjacent mouse stroma.

Viral integration frequently occurs at “fragile sites;” these genomic regions prone to replication stress and DNA breakage which would create susceptible sites for integration events^13^. If replication errors remain unresolved, chromosomal instability further increases opportunities for integration. We previously demonstrated that fibroblasts regulate both DNA repair pathways and numerous histones at the RNA level^10^. Here we demonstrate that the presence of fibroblasts supported overall DNA integrity in our HPV+ models. Co-culture with fibroblasts decreased DNA strand breaks, at least in portion by the regulation of replication fork dynamics and the expression of DNA damage repair factors^10^.

Future studies will aim to develop patient-derived cell lines using this co-culture approach. This methodology has already proven effective for generating episomal cell lines from primary keratinocytes and CIN lesions, suggesting that a similar strategy could facilitate both the establishment and episomal maintenance of lines derived from cancerous lesions. Suggested fibroblast plating densities are further defined in the methods section. Establishing robust episomal HPV+ OPC models will provide a critical resource for advancing mechanistic insights and translational research.

## Materials and Methods

### Cell Culture

UMSCC104 (Millipore) and UMSCC152 (ATCC) cells were grown in Eagle’s minimum essential medium (EMEM) (Invitrogen) supplemented with nonessential amino acids (NEAA) (Gibco) and 20% fetal bovine serum (FBS) (Gibco) and were grown in the presence or absence of inactivated fibroblast feeder cells (described below). HN4 (generous gift from Dr. Hisashi Harada and FTR fingerprinted) and UMSCC47 (Millipore) cells were grown in were grown in Dulbecco’s modified Eagle’s medium (DMEM) (Invitrogen; Carlsbad, CA, USA) supplemented with 10% FBS. N/Tert-1 and N/Tert-1+HPV16 cells were maintained in keratinocyte-serum free medium (K-SFM; Invitrogen); +HPV16 cells were supplemented with 150µg/ml G418 (Fisher Scientific). HFK^+^HPV16 have been previously described, were grown in Dermalife-K complete media (Lifeline Technology)^18^. All cells were maintained at 37°C in a 5% CO_2_–95% air atmosphere, passaged every 3 or 4 days, and routinely monitored for mycoplasma (Sigma, MP0035).

### Culture and mitomycin C (MMC) inactivation of 3T3-J2 mouse fibroblast feeder cells

3T3-J2 immortalized mouse fibroblasts (J2) were grown in DMEM (Gibco) and supplemented with 10% FBS. 70-80% confluent plates were treated with 4µg/ml of MMC in DMSO (Cell Signaling Technology) for 4-6 hours at 37°C, as previously described^10^. For co-culture conditions, every 2-3 days, 100-mm plates were supplemented with 1x10^6^ J2, 6-well plates were supplemented with 1x10^5^ J2, or 24-well plates were supplemented with 1x10^4^ J2. For long term cultures, residual J2s were washed off with 1x PBS and new J2 were continually supplemented every 2-3 days.

### Immunoblotting

Specified cells were washed with 1X PBS, trypsinized, lysed, and ran as previously described^10^. Membranes were blocked with Odyssey T-PBS blocking buffer at room temperature for at least 1 hour and probed with indicated primary antibody diluted in Odyssey blocking buffer. Membranes were washed with PBS supplemented with 0.1% Tween (PBS-Tween) and probed with the indicated Odyssey secondary antibody 1:10,000 (goat anti-mouse IRdye 800CW or goat anti-rabbit IRdye 680CW) diluted in Odyssey blocking buffer. Membranes were washed three times with PBS-Tween and an additional wash with 1X PBS. Infrared imaging of the blot was performed using the Odyssey CLx Li-Cor imaging system. The following primary antibodies were used for immunoblotting in this study at 1:500: E2^25^, GAPDH 1:10,000 (Santa Cruz, sc-47724).

### Reproducibility, research integrity, and statistical analysis

All experiments were carried out at least in triplicate in all of the cell lines indicated. All cell lines were bought directly from sources indicated, or typed via cell line authentication services. All images shown are representatives from triplicate experiments. Quantification is presented as mean +/- standard error (SE). Student’s t-test was used to determine significance: * p<0.05, **p<0.01, ***p<0.001

## Acknowledgements

This work was supported by the VCU Philips Institute for Oral Health Research, the OneVCU Research Quest Fund, and the National Cancer Institute-designated Massey Cancer Center grant P30 CA016059.

